# Combined nelfinavir and cisplatin treatment promotes inflammatory signaling and immunogenic cell death in ovarian cancer cells

**DOI:** 10.64898/2026.11.16.767109

**Authors:** Benjamin N. Forgie, Sarah T. Ferrier, Alicia A. Goyeneche, Alexandra Bartolomucci, Desiree Marno, Rewati Prakash, Abu Shadat M. Noman, Edith Zorychta, Lucy Gilbert, Julia V. Burnier, Carlos M. Telleria

## Abstract

Platinum (Pt)-based chemotherapy remains central to ovarian cancer (OC) treatment, but recurrent disease frequently develops Pt resistance and responds poorly to immune checkpoint inhibitors. Immunogenic cell death (ICD) links tumor cell killing to antitumor immunity; however, cisplatin (CDDP), widely used in OC, is an incomplete ICD inducer. We previously demonstrated that the HIV protease inhibitor nelfinavir (NFV) enhances CDDP-induced OC cell death and promotes a lytic, pyroptosis-associated phenotype. Here, we investigated whether NFV enhances the immunogenic consequences of CDDP treatment. RNA-seq analysis of OC cells treated with CDDP, NFV, or their combination revealed enrichment of interferon (IFN)-associated, cytoplasmic DNA-sensing, inflammatory chemokine, and antigen-presentation pathways following NFV-containing treatments. NFV alone and CDDP/NFV reduced mature Lamin A abundance and increased extranuclear double-stranded DNA accumulation. Cytosolic fractionation and droplet digital PCR demonstrated increased cytosolic *TP53* DNA, particularly following combined treatment. CDDP/NFV increased STING and TBK1 phosphorylation, supporting engagement of the cGAS–STING–TBK1 signaling pathway. The combination enhanced release of inflammatory and ICD-associated mediators, including HMGB1, IL-18, and IFN-β. Importantly, vaccination with CDDP/NFV-treated murine ID8 OC cells protected immunocompetent female mice against subsequent live ID8 cell challenge, reducing tumor incidence and burden. In contrast, ID8 cells killed by freeze-thaw-induced accidental necrosis provided no comparable protection. Collectively, these findings indicate that NFV enhances the immunogenic consequences of CDDP-induced OC cell death, potentially involving cytoplasmic DNA accumulation and STING–TBK1 signaling, and promotes protective antitumor immunity *in vivo*. These results support further investigation of NFV as a repurposed partner for Pt-based chemotherapy and its potential combination with immunotherapy in OC.

## 1. Introduction

Ovarian cancer (OC) remains the most lethal gynecologic malignancy [1], with most patients diagnosed at advanced stages and a poor 5-year survival for disseminated disease [2]. Although immune checkpoint inhibitors (ICIs) have transformed the treatment of several malignancies, including melanoma [3], non-small cell lung cancer [4], endometrial and cervical cancers [5], and renal cell carcinoma [6], their efficacy in OC remains limited, with response rates of approximately 10-15% [7]. This limitation does not necessarily reflect an absence of tumor-infiltrating immune cells, as increased CD8+ T-cell infiltration is associated with improved clinical outcomes [8]. Nevertheless, ICIs, alone or combined with chemotherapy, provide limited benefit in unselected OC populations [7,9,10], potentially reflecting features of the tumor immune microenvironment (TIME), including immune exclusion [11], suppressive myeloid and regulatory T-cell populations [12], low tumor mutational burden [13], impaired antigen presentation [14], or inconsistent predictive value of programmed death-ligand 1 (PD-L1) expression [15]. These findings highlight the need for therapeutic strategies that enhance tumor immunogenicity and inflammatory signaling beyond direct tumor cell killing.

Immunogenic cell death (ICD) represents one mechanism through which therapies can promote anti-tumor immunity [16]. This form of regulated cell death couples tumor cell killing with the exposure or release of immune-stimulatory signals, including damage-associated molecular patterns (DAMPs) [17], inflammatory cytokines and chemokines, type I interferons (IFNs) [18], and other mediators that support adjuvancy, antigen presentation, and adaptive immune activation [19]. However, not all cytotoxic agents induce ICD. Although oxaliplatin (OXP) is a recognized ICD-inducing platinum (Pt) derivative [20], cisplatin (CDDP) and carboplatin (CP), the Pt drugs most widely used in OC treatment) [21], are comparatively weak or incomplete ICD inducers [22–24]. Thus, strategies that redirect Pt-induced OC cell death toward a more immunogenic phenotype, characterized by coordinated adjuvant and antigenic signals, may enhance the immune-relevant consequences of conventional chemotherapy.

Cytoplasmic DNA sensing links therapy-induced cellular damage to inflammatory signaling [25]. DNA damage [26], nuclear envelope disruption [26], micronuclei formation [26], and mitochondrial stress [27] can promote cytoplasmic accumulation of double-stranded (ds) DNA, activating cyclic GMP–AMP synthase (cGAS) and its downstream effector, stimulator of interferon genes (STING) [25–27]. STING activation recruits TANK-binding kinase 1 (TBK1), which phosphorylates interferon regulatory factor 3 (IRF3), inducing type I IFN production and inflammatory transcriptional programs [28]. These responses can promote antitumor immunity through dendritic cell activation, antigen presentation, effector T-cell recruitment, and natural killer cell activation [29,30]. Accordingly, pharmacologic STING agonists, alone or in combination with ICIs, are investigated clinically to enhance tumor inflammation and potentially improve immunotherapy responses [29,31]. Although STING and type I IFN signaling can have context-dependent effects, cytoplasmic DNA sensing remains an important mechanism linking cellular damage to immune-relevant inflammation [26].

We previously demonstrated that the HIV protease inhibitor nelfinavir (NFV) synergizes with CDDP to induce intracellular stress and OC cell death, promoting a lytic, pyroptosis-associated phenotype [32]. NFV has also been reported to interfere with Lamin A maturation, potentially compromising nuclear-envelope integrity [33,34]. Given the role of nuclear envelope disruption in cytoplasmic DNA accumulation and cGAS-STING activation [26], we hypothesized that NFV could enhance the otherwise poorly inflammatory and immunogenic consequences of CDDP-induced cell death [22]. We therefore investigated whether NFV redirects CDDP-induced cell death towards a more immunogenic phenotype by examining cytoplasmic DNA accumulation, cGAS-STING pathway activation, inflammatory transcriptional reprogramming, and the release of soluble inflammatory mediators. Finally, we evaluated the functional immunogenic consequences of this drug combination on the death of OC cells in a prophylactic vaccination model.

## 2. Materials and methods

### 2.1. Cell culture and reagents

Human IGROV-1/CP, PEO4, and TOV21G OC cells and murine ID8 OC cells were used. PEO4, IGROV-1/CP, TOV21G, and ID8 cells were provided by Dr. Taniguchi (Fred Hutchinson Cancer Center, University of Washington, Seattle, WA, USA), Dr. Stephen Howell (University of California, San Diego, CA, USA), Dr. Mes-Masson (Centre de recherche du Centre Hospitalier de l’Université de Montréal [CHUM], Montréal, QC, Canada), and Dr. Vanderhyden (University of Ottawa, Ottawa, ON, Canada), respectively. All cell lines were maintained in RPMI 1640 medium (Mediatech, Manassas, VA, USA) supplemented with 10% fetal bovine serum (FBS; Corning Inc., Corning, NY, USA), 0.01 mg/mL human insulin (Roche, Indianapolis, IN, USA), 10 mM HEPES (Corning), 100 IU/mL penicillin (Mediatech), 100 µg/mL streptomycin (Mediatech), 2 mM L-alanyl-L-glutamine (GlutaGro™, Corning), and 1 mM sodium pyruvate (Corning). Cells were cultured at 37 °C in a humidified incubator with 5% CO. Cis-diamminedichloroplatinum II (cisplatin [CDDP]; Sigma Chemical Co., St. Louis, MO, USA) was dissolved in 0.9% sodium chloride. Nelfinavir mesylate hydrate (NFV; Sigma) was dissolved in dimethyl sulfoxide (DMSO; Sigma). Vehicle controls received equivalent solvent concentrations, with final DMSO concentrations not exceeding 0.1% (v/v).

### 2.2. Drug treatments

For *in vitro* experiments, cells received vehicle, CDDP, NFV, or CDDP/NFV. Vehicle controls were exposed to 0.9% sodium chloride for 3 h, followed by complete medium containing DMSO for 72 h. CDDP (10 µM) was administered for 3 h, followed by drug-free complete medium for 72 h. NFV-treated cells received vehicle for 3 h, followed by 20 µM NFV for 72 h. Combination-treated cells received 10 µM CDDP for 3 h, followed by 20 µM NFV in complete medium for 72 h.

### 2.3. RNA sequencing and Reactome pathway analysis

RNA-seq data from vehicle-, NFV-, and CDDP/NFV-treated IGROV-1/CP cells were reanalyzed for immune- and inflammation-related transcriptional programs, as previously described [32]. Differential expression analysis was performed using DESeq2 (v1.46.0), followed by preranked gene set enrichment analysis (GSEA) using fgsea (v1.32.4) and Reactome gene sets (MSigDB v2025.1.Hs). Selected immune- and inflammation-associated pathways were visualized by enrichment and dot plots. Variance-stabilizing transformed expression values were row-scaled to Z-scores for heatmap visualization of selected immune-related genes.

### 2.4. Immunoblotting

Following treatment, OC cells were scraped into ice-cold 1X PBS, pelleted by centrifugation, and stored at −80 °C. Cells were lysed in NP-40 lysis buffer containing protease inhibitors for 30 min on ice. Lysates were clarified by centrifugation (12,000 × *g*, 15 min, 4 °C), and protein concentrations were determined using the Pierce BCA Protein Assay Kit (Thermo Fisher Scientific, Waltham, MA, USA). Twenty-five µg of total protein per lane was separated on 12% SDS-polyacrylamide gels prepared using the TGX Stain-Free FastCast Acrylamide Kit (Bio-Rad Laboratories, Hercules, CA, USA) at 200 V for 35 min and transferred onto PVDF membranes using the Trans-Blot Turbo Transfer System (1.3 A, 7 min; Bio-Rad). Membranes were blocked with 5% non-fat milk in TBS-T for 1 h at room temperature (RT), followed by overnight incubation at 4 °C with primary antibodies (1:1000) diluted in either 5% non-fat milk or 5% bovine serum albumin (BSA; Sigma) in TBS-T. Primary antibodies included Lamin A (sc-518013, Santa Cruz Biotechnology, Dallas, TX, USA), phospho-STING (ser366; #19781), and phospho-TBK1 (ser172; #5483; Cell Signaling Technologies, Danvers, MA, USA), and β-actin (A5441, Sigma). After washing with TBS-T, membranes were incubated for 1 h at RT with HRP-conjugated goat anti-rabbit IgG (H+L; 1706515) or goat anti-mouse IgG (H+L; 1706516; Bio-Rad). Bands were visualized using Clarity Western Blot ECL Substrate (Bio-Rad) and imaged with a ChemiDoc Touch Imaging System (Bio-Rad). Experiments were independently performed at least twice, and representative immunoblots are shown.

### 2.5. Immunofluorescence

Cells were seeded in 8-well Nunc™ Lab-Tek™ II CC2™ Chamber Slide Systems (Thermo Fisher) and treated as described above. After three PBS washes, cells were fixed with 4% paraformaldehyde for 20 min at RT, permeabilized with 0.1% Triton X-100 for 5 min, and blocked with 1% BSA for 20 min at RT, with three PBS washes after permeabilization. Cells were incubated overnight at 4 °C with an anti-dsDNA antibody (sc-58749, Santa Cruz) diluted 1:100 in 1% BSA, followed by Alexa Fluor™ 488-conjugated goat anti-mouse IgG (H+L) cross-adsorbed secondary antibody (A32723; Thermo Fisher) diluted 1:1000 for 1 h at RT. Nuclei were counterstained with 300 nM DAPI for 10 min at RT. After PBS washes, images were acquired using a Cytation 3 Cell Imaging Multi-Mode Reader (BioTek Instruments, Winooski, VT, USA). Experiments were independently performed at least twice. Representative images acquired under identical exposure settings are shown; brightness and contrast were adjusted uniformly across treatment conditions within each cell line, where applicable.

### 2.6. Cytoplasmic DNA isolation

Following treatment, cells were resuspended in ice-cold lysis buffer (10 mM Tris-HCl, 10 mM NaCl, 3 mM MgCl, 300 mM sucrose, and 0.05% NP-40) and incubated for 20 min on ice. Samples were centrifuged at 500 × *g* for 5 min at 4 °C to pellet bulk nuclei, followed by centrifugation of the supernatants at 10,000 × *g* for 5 min at 4 °C. Cytosolic fractions were collected and stored at −80 °C. DNA was extracted using the EZ2 automated and EZ2 ccfDNA Kit (cat. no. 954854; QIAGEN, Hilden, Germany), according to the manufacturer’s instructions, and stored at −20 °C until ddPCR analysis.

### 2.7. Droplet digital PCR analysis

Primers, hydrolysis probes, and synthetic DNA controls (gBlock gene fragments) were designed and synthesized by Integrated DNA Technologies (IDT, Coralville, IA, USA). Droplet digital PCR (ddPCR) reactions were prepared according to the manufacturer’s instructions using 10 μL of 2 × ddPCR Supermix for Probes (No UTP; Bio-Rad), 900 nM of each forward and reverse primer, 250 nM FAM-or HEX-labelled Affinity Plus qPCR probes, 2 μL of DNA template, and nuclease-free water (IDT). Positive and negative gBlock controls and no-template controls were included. Reaction mixtures were loaded into droplet generation cartridges with 70 μL of Droplet Generation Oil (Bio-Rad). Droplets were generated using the QX200 Droplet Generator (Bio-Rad) and transferred to a 96-well PCR plate. Thermal cycling consisted of 95 °C for 10 min; 50 cycles of 95 °C for 30 s, primer-specific annealing at 54–62 °C for 1 min, and 72 °C for 30 s; and 98 °C for 5 min with a 2 °C/s ramp rate. Annealing temperatures were optimized for each primer/probe set. Droplets were analyzed using the QX200 Droplet Reader (Bio-Rad), and data were processed with QuantaSoft software. Cytosolic *TP53* copy numbers were normalized to the cell numbers and expressed as copies per 100,000 cells. Three independent biological experiments were performed, with technical duplicates for each sample. Reactions yielding fewer than 10,000 droplets were excluded, and samples with ≤ 2 mutant-positive droplets were considered negative. Cell-specific *TP53* primer/probe sets targeted the respective *TP53* mutation loci: *TP53*c377 for IGROV-1/CP cells and *TP53*c731 for PEO4 cells. The primers and probes used are as follows: TP53 c377 Forward: TCAACTGTCTCTCCTTCCTCTT; TP53 c377 Reverse: TGTGGAATCAACCCACAGC; TP53 c731 Forward: CTGTACCACCATCCACTACAAC; TP53 c731 Reverse: TCCTGACCTGGAGTCTTCC; TP53 c377A>G mutant probe: /56-FAM/ACA G+T+G +CTC +CC/3IABkFQ/; TP53 c377 WT probe: /5HEX/CAG +T + A +C TC+C + CC/3IABkFQ/; TP53 c731G>A mutant probe: /56-FAM/ATG +G+A+C GGC AT+G A/3IABkFQ/; TP53 c731 WT probe: /5HEX/ATG G+G+C GGC AT+G A/3IABkFQ/.

### 2.8. Measurement of soluble inflammatory mediators

Culture supernatants were collected after treatment and analyzed using commercial immunoassays according to the manufacturers’ instructions. Human IFN-β was measured using the Human IFN beta ELISA Kit (ab278127; Abcam, Cambridge, UK), HMGB1 using the Lumit® HMGB1 Human/Mouse Immunoassay (W6110; Promega Corporation, Madison, WI, USA), and total IL-18 using the Human Total IL-18 DuoSet ELISA (DY318-05; R&D Systems, Minneapolis, MN, USA). ELISA absorbance was measured at 450 nm using the Cytation 3 Cell Imaging Multi-Mode Reader (BioTek Instruments), and concentrations were calculated from standard curves. HMGB1 luminescence was also measured using the Cytation 3 Cell Imaging Multi-Mode Reader (BioTek Instruments). Data are expressed as relative luminescence units or supernatant concentrations, as indicated. Three independent biological experiments were performed per condition, and technical replicates were averaged before statistical analysis.

### 2.9. Prophylactic vaccination studies

Murine ID8 OC cells were used for the prophylactic vaccination assay. For vaccine preparation, cells were treated with 10 µM CDDP for 3 h, followed by 20 µM NFV for 72 h. Floating and adherent cells were collected, and cell number and viability were assessed by trypan blue exclusion. Preparations containing approximately 80% dying/death cells were used for vaccination. As a non-ICD control, ID8 cells underwent accidental necrosis through two liquid nitrogen freeze-thaw cycles. Both preparations were collected and resuspended in their respective culture supernatants to preserve soluble factors released during cell death. Female C57BL/6 mice (20-22 g; Cat. 027C57BL/6 Mouse, Charles River Labs, St. Constant, QC, Canada) were randomly assigned to three groups: PBS control (n = 10), CDDP + NFV vaccine (n = 10), and freeze-thaw control (n = 6). On days 1 and 7, mice received subcutaneously (s.c.) injections in the right flank of 5 × 10 ID8 cells in 100 µL. CDDP/NFV-treated preparations contained approximately 80% dying/death cells, whereas freeze–thaw preparations contained 100% dead cells. PBS controls received 100 µL PBS. On day 14, all mice were rechallenged s.c. in the opposite flank with 1 × 10 untreated ID8 cells with > 95% viability. Mice were monitored every 2 days for palpable tumors. Tumor-free survival was defined as the interval from live ID8 rechallenge to first palpable tumor detection; mice without palpable tumors at day 80 were right-censored. Tumor dimensions were measured using an electronic caliper (0.01 mm resolution), and volumes were calculated as V = (L × W²)/2, where L and W represent the longest tumor dimension and perpendicular width, respectively, as previously reported [35]. Animal procedures were approved under protocol MCGL-10174 and conducted in accordance with institutional animal-care guidelines.

### 2.10. Statistics

Statistical analyses were performed using GraphPad Prism (v10.6.1) and R. Quantitative *in vitro* data are presented as mean ± SD from three independent biological experiments, unless otherwise indicated. Two-group comparisons used unpaired two-tailed Student’s *t*-tests, whereas comparisons among three or more groups used one-way ANOVA followed by Tukey’s multiple-comparisons test when parametric assumptions were met. Tumor-free survival was estimated using the Kaplan–Meier method and compared between each vaccine group and PBS controls using the Mantel-Cox log-rank test. Time to event was defined as days from live ID8 rechallenge to first palpable tumor detection; tumor-free mice at day 80 were right-censored. Endpoint tumor volumes were compared using the Kruskal–Wallis test followed by Dunn’s multiple-comparisons test. Two-sided *p* values < 0.05 were considered statistically significant. Differential gene expression was analyzed using DESeq2, with genes meeting an FDR-adjusted *p* value < 0.05 and absolute log fold change ≥ 1 considered differentially expressed. Reactome pathway enrichment was performed using fgsea with genes ranked by the DESeq2 Wald statistic; pathways with FDR-adjusted *p* values < 0.05 were considered significantly enriched. Heatmaps were used for visualization only and were not subjected to separate statistical testing.

## 3. Results

### 3.1. NFV-containing treatments induce interferon-associated inflammatory and antigen presentation transcriptional programs

To determine whether NFV-containing treatments induce immune-related transcriptional reprogramming, we analyzed RNA-seq data from IGROV-1/CP cells treated with vehicle, CDDP, NFV, or CDDP/NFV (Fig. 1A). Gene set enrichment analysis of the three treatment-versus-vehicle controls revealed that NFV and CDDP/NFV induced the strongest positive enrichment of selected immune- and inflammation-associated Reactome pathways, whereas CDDP alone generally elicited weaker enrichment. The pathways included cytoplasmic DNA sensing, type I IFN signaling, modulation of host responses by IFN-stimulated genes, TNF signaling, class I MHC antigen presentation, MyD88-independent TLR4 signaling, and TLR9 signaling (Fig. 1B).

**Fig. 1.**
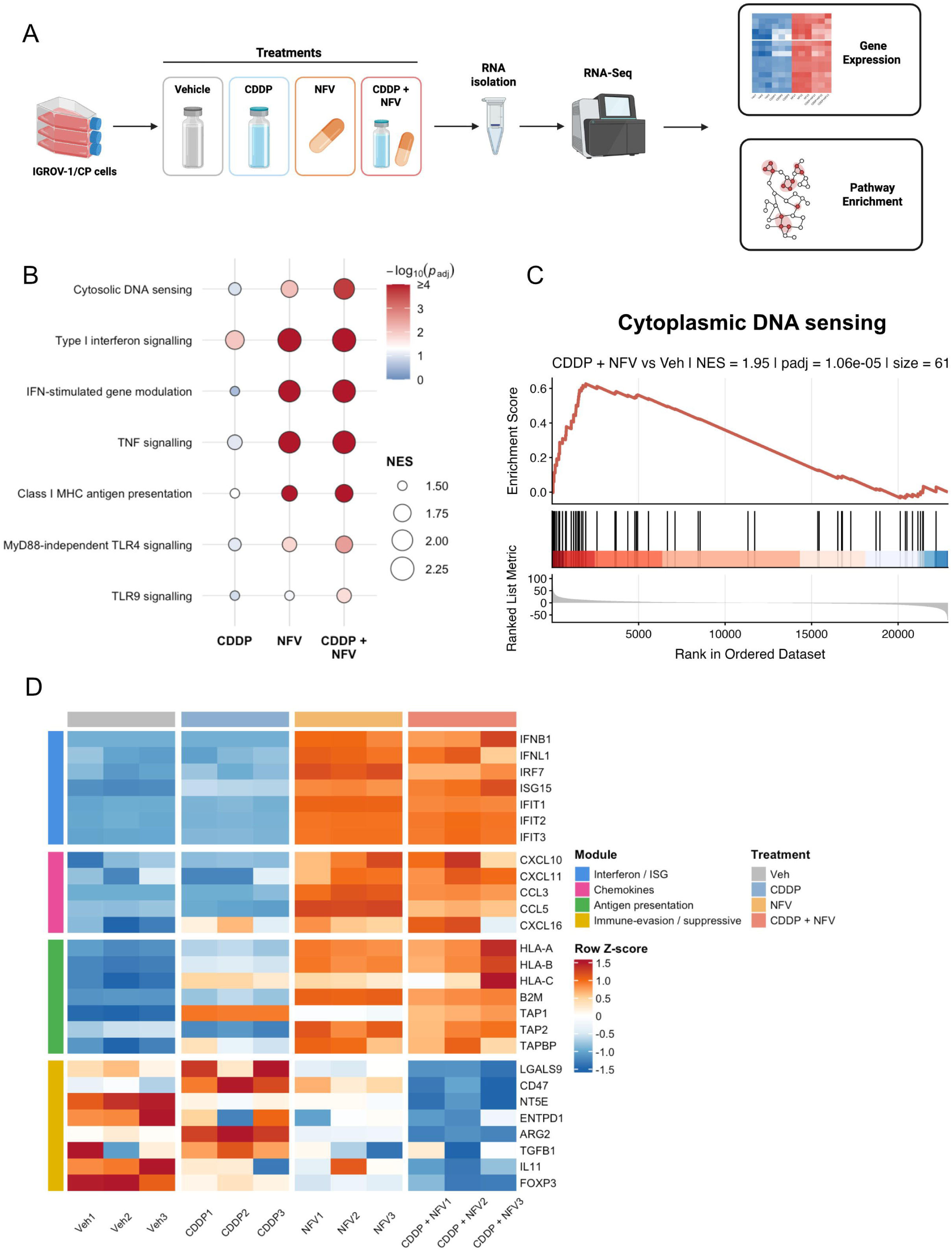
NFV-containing treatments induce IFN-related, IFN-associated, and antigen-presentation transcriptional programs in IGROV-1/CP cells. A) RNA-seq workflow. IGROV-1/CP cells were treated with vehicle, CDDP, NFV, or CDDP + NFV and collected at 24 h. Three independent biological replicates were analyzed per condition. B) Gene set enrichment analysis of selected immune- and IFN-related Reactome pathways in treated versus vehicle-control cells. Dot size represents the normalized enrichment score (NES), and color indicates −log of the FDR-adjusted *p* value (capped at 4). C) Running enrichment plot for the Reactome cytosolic DNA-sensing pathway in CDDP + NFV-treated cells relative to vehicle control cells. Vertical ticks indicate pathway-associated genes in the ranked list; the lower track shows the DESeq2 Wald statistic. NES, FDR-adjusted *p* value, and gene-set size are indicated above the plot. D) Heatmap of selected genes grouped into interferon/interferon stimulated genes (ISG), chemokine, class I antigen-presentation, and immune-evasion/suppressive modules. Variance-stabilizing transformed expression values are displayed as row Z-scores, capped at ±1.5. Columns represent individual biological replicates; colored annotations indicate treatment and gene module.

Among these pathways, genes associated with cytoplasmic DNA sensing showed enrichment toward the top of the ranked gene list in CDDP/NFV-treated cells relative to vehicle controls (NES = 1.95, FDR-adjusted *p* = 1.06 × 10; Fig. 1C). This finding provided a transcriptional rationale for subsequently investigating cytoplasmic DNA accumulation and downstream DNA-sensing signaling in response to treatment.

Consistent with the pathway-level findings, NFV and CDDP/NFV increased expression of selected IFN-associated transcripts, including *IFNB1*, *IFNL1*, *IRF7*, *ISG15*, *IFIT1*, *IFIT2*, and *IFIT3* (Fig. 1D). Both treatments also increased the expression of inflammatory chemokines, (*CXCL10, CXCL11*, *CCL3*, *CCL5*, and *CXCL16*), and genes involved in class I antigen processing and presentation (*HLA-A*, *HLA-B*, *HLA-C*, *B2M*, *TAP1*, *TAP2,* and *TAPBP*) (Fig. 1D). Conversely, CDDP/NFV generally reduced the expression of selected immunomodulatory or immunosuppressive transcripts, including *LGALS9*, *CD47*, *NT5E*, *ENTPD1*, *ARG2*, *TGFB1*, *IL11*, and *FOXP3* (Fig. 1D). Overall, these findings identify NFV as a major contributor to the inflammatory transcriptional response and indicated that both NFV and CDDP/NFV promote IFN-associated, chemokine-rich, and antigen-presentation expression profiles in IGROV-1/CP cells.

### 3.2 NFV drives cytoplasmic self-dsDNA accumulation and activation of STING signaling

Given the enrichment of cytoplasmic DNA-sensing and IFN-associated pathways following NFV-containing treatments, we next examined whether these transcriptional changes were accompanied by extranuclear dsDNA accumulation and activation of STING-associated signaling. NFV has previously been reported to interfere with Lamin A maturation [33,34], and Lamin A is an integral structural component of the nuclear lamina [36]. This raised the possibility that NFV-induced alterations in Lamin A may contribute to the accumulation of self-DNA outside the nucleus. In PEO4 and IGROV-1/CP cells, NFV-containing treatments reduced mature Lamin A abundance compared with vehicle and CDDP alone (Fig. 2A). A pronounced reduction was also observed following CDDP/NFV treatment in TOV21G cells (Supplementary Fig. S1A). These findings are consistent with treatment-associated alterations in Lamin A abundance or maturation but do not, by themselves, establish disruption of nuclear-envelope integrity.

**Fig. 2.**
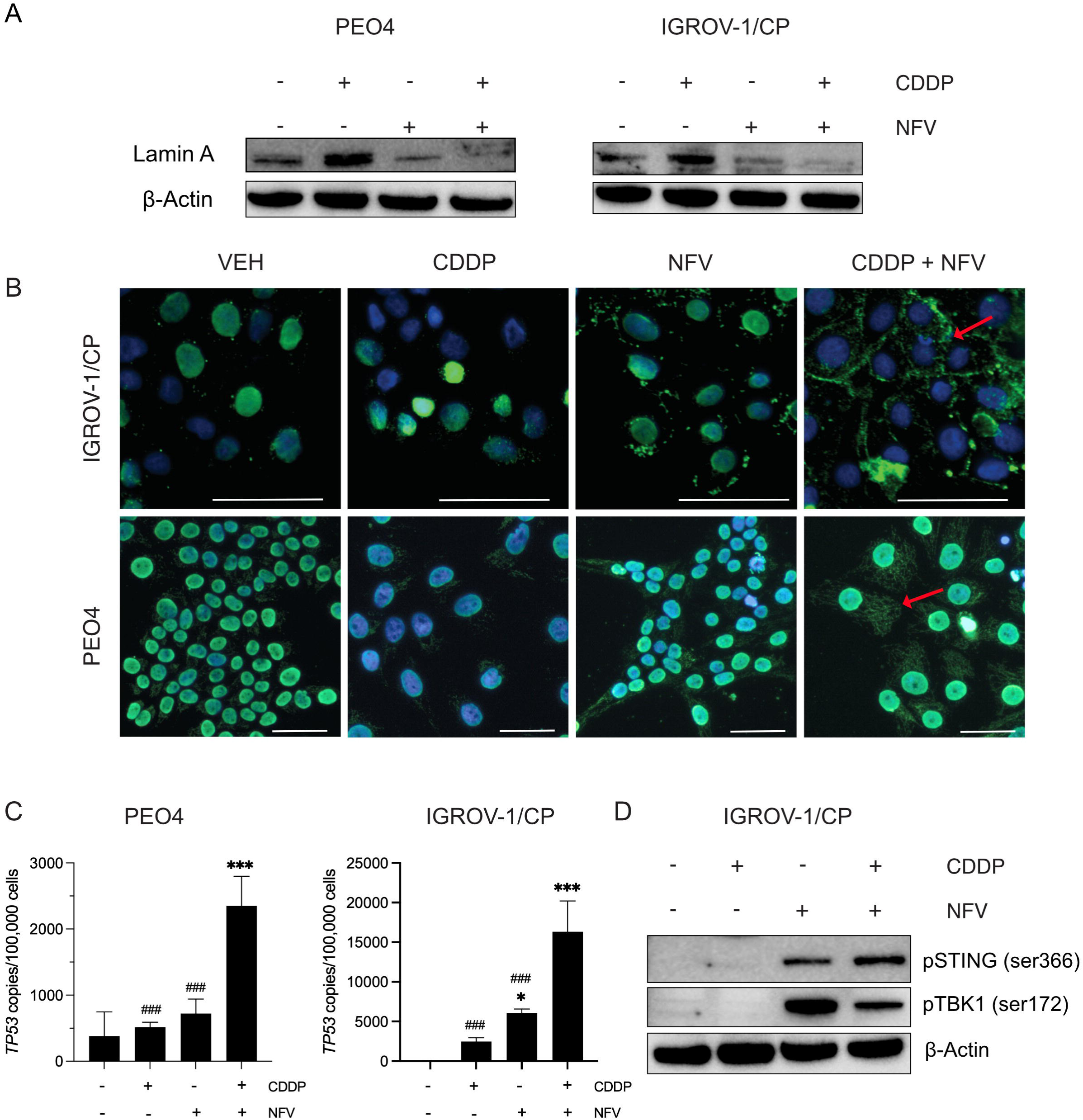
NFV promotes cytoplasmic DNA accumulation and STING pathway activation in ovarian cancer cells. A) Representative immunoblots of Lamin A in PEO4 and IGROV-1/CP cells following the indicated treatments; β-Actin served as a loading control. B) Representative immunofluorescence images showing dsDNA (green) and DAPI-stained nuclei (blue); red arrows indicate representative extranuclear dsDNA. Scale bars: 50 µm. C) Cytosolic *TP53* DNA copies quantified by ddPCR and normalized to 100,000 cells. Data are mean ± SD from three independent biological experiments, each analyzed in technical duplicate. D) Representative immunoblots of phosphorylated STING (ser366) and TBK1 (ser172) in IGROV-1/CP cells; β-Actin served as a loading control. Immunoblotting and immunofluorescence were performed independently at least twice. Statistical analysis in C used one-way ANOVA followed by Tukey’s multiple-comparisons test. \**p* < 0.05 and \*\*\**p* < 0.001 versus vehicle; ###*p* < 0.001 versus CDDP + NFV.

Immunofluorescence analysis revealed predominantly nuclear dsDNA staining in vehicle-treated cells, whereas NFV- and CDDP/NFV-treated cells exhibited increased extranuclear dsDNA signals in both IGROV-1/CP and PEO4 cells (Fig. 2B). CDDP alone generally produced less extranuclear dsDNA staining than either NFV-containing treatment. To quantify cytoplasmic DNA accumulation, we isolated cytosolic fractions and performed ddPCR using cell-line-specific *TP53* detection. CDDP/NFV treatment increased cytosolic *TP53* DNA copy numbers in both PEO4 and IGROV-1/CP cells relative to vehicle controls (Fig. 2C). NFV alone also increased cytosolic *TP53* copies in IGROV-1/CP cells, whereas the increase was modest and did not reach statistical significance in PEO4 cells.

Finally, we examined phosphorylation of STING and its downstream kinase TBK1. In IGROV-1/CP cells, NFV and CDDP/NFV increased STING and TBK1 phosphorylation relative to vehicle and CDDP treatments (Fig. 2D). TOV21G cells also displayed STING and TBK1 phosphorylation, with the strongest TBK1 phosphorylation observed following CDDP/NFV treatment (Supplementary Fig. S1B). In contrast, phosphorylation of STING-pathway components was variable and not consistently detected in PEO4 cells (data not shown). Collectively, these findings associate NFV-containing treatments with reduced Lamin A abundance and increased extranuclear DNA accumulation in OC cells, accompanied by STING-TBK1 pathway activation most clearly in IGROV-1/CP and TOV21G cells, but not consistently detected in PEO4 cells.

### 3.3 Death of OC cells caused by CDDP/NFV treatment is associated with release of DAMPs and cytokines

After observing cytoplasmic dsDNA accumulation and STING-pathway activation in OC cells treated with NFV alone or in combination with CDDP, we investigated whether these changes were accompanied by the release of soluble mediators associated with inflammation and ICD. We first measured extracellular levels of high-mobility group box 1 (HMGB1), a non-histone DNA-binding and repair protein that functions as a DAMP associated with ICD [37]. In PEO4 cells, HMGB1 release was highest following CDDP/NFV treatment, whereas either drug alone produced more modest increases (Fig. 3A). In IGROV-1/CP cells, both CDDP and NFV increased extracellular HMGB1 relative to vehicle controls, with the highest levels again observed following combination treatment (Fig. 3A).

**Fig. 3.**
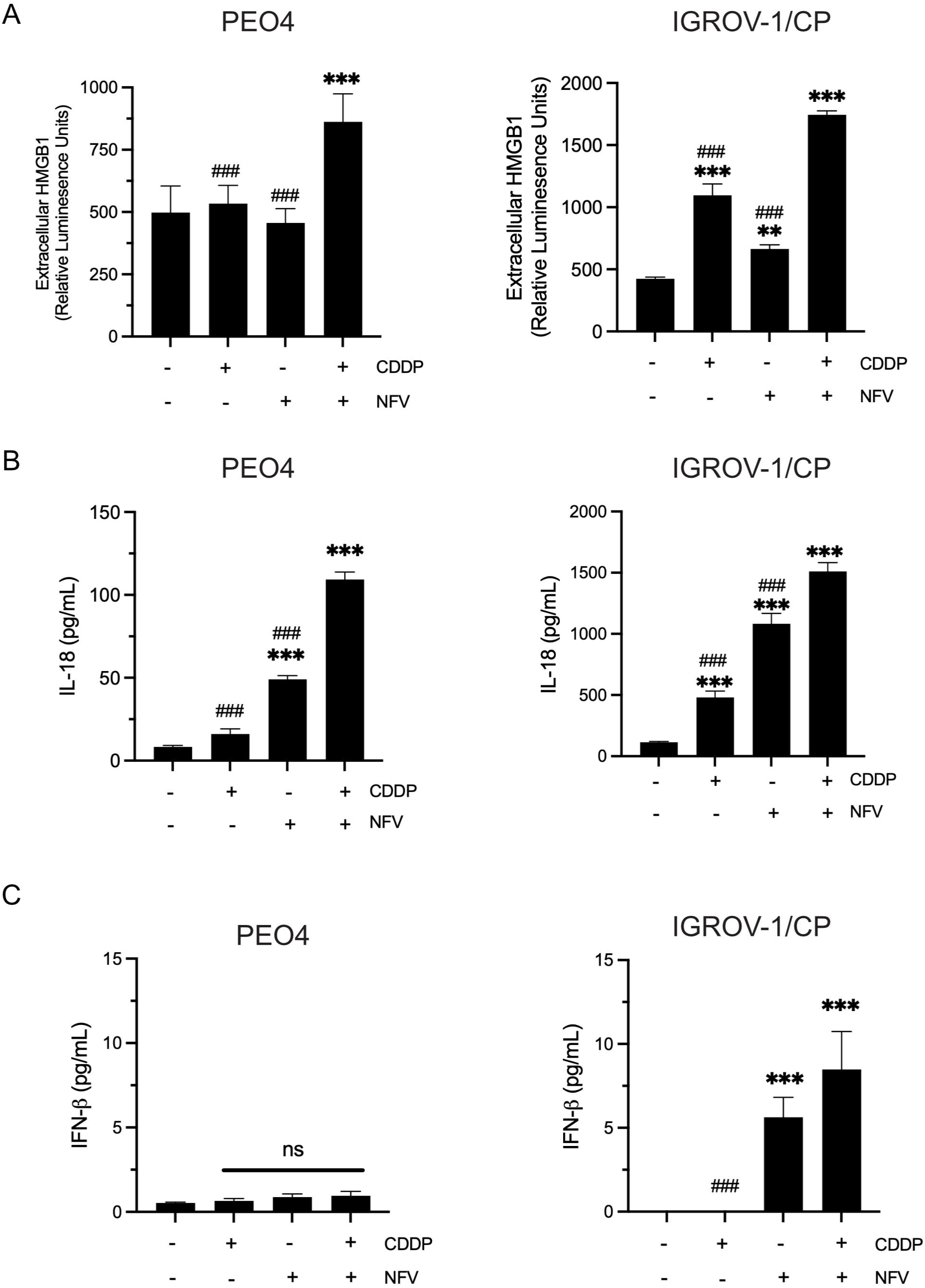
NFV-containing treatments promote release of soluble mediators associated with inflammatory and ICD. A-C) Extracellular HMGB1, IL18, and INF-β levels in culture supernatants from PEO4 and IGROV-1/CP cells treated with vehicle, CDDP, NFV, or CDDP + NFV. HMGB1 release is expressed in relative luminescence units. Data are the mean ± SD from three independent biological experiments. Statistical significance was determined by one-way ANOVA followed by Tukey’s multiple-comparisons test. \*\**p* < 0.01 and \*\*\**p* < 0.001 versus vehicle; ###*p* < 0.001 versus CDDP + NFV; ns, not significant.

We next measured IL-18, an inflammatory cytokine constitutively produced and stored by many epithelial cells [38–40] and associated with immunostimulatory forms of cell death [41]. NFV increased IL-18 secretion, which CDDP/NFV treatment further enhanced in both PEO4 and IGROV-1/CP cells (Fig. 3B). IGROV-1/CP cells showed a particularly pronounced response, with both single-agent treatments increasing IL-18 release and the combination producing the highest extracellular levels (Fig. 3B).

Finally, we assessed IFN-β release as a downstream functional readout of cytoplasmic self-dsDNA sensing and STING-associated inflammatory signaling [42]. Consistent with the STING and TBK1 phosphorylation detected by immunoblotting, both NFV alone and CDDP/NFV increased IFN-β secretion in IGROV-1/CP cells (Fig. 3 C). In contrast, IFN-β levels in PEO4 cells were very low and did not differ among treatment groups, consistent with the absence of robust STING-pathway activation in these cells (data not shown).

### 3.4 Vaccination with CDDP/NFV-treated ID8 cells reduces tumor incidence and tumor burden

To determine whether CDDP/NFV-induced cell death was functionally immunogenic, beyond its association with the release of ICD-associated biomarkers such as HMGB1 and IL-18, we employed a prophylactic vaccination model in immunocompetent C57BL/6 mice. Wild-type murine ID8 OC cells were treated *ex vivo* with CDDP/NFV to generate a vaccine preparation containing approximately 80% dying or dead cells. Female mice were vaccinated on days 1 and 7 and rechallenged on day 14, on the opposite flank, with live untreated ID8 cells (Fig. 4A).

**Fig. 4.**
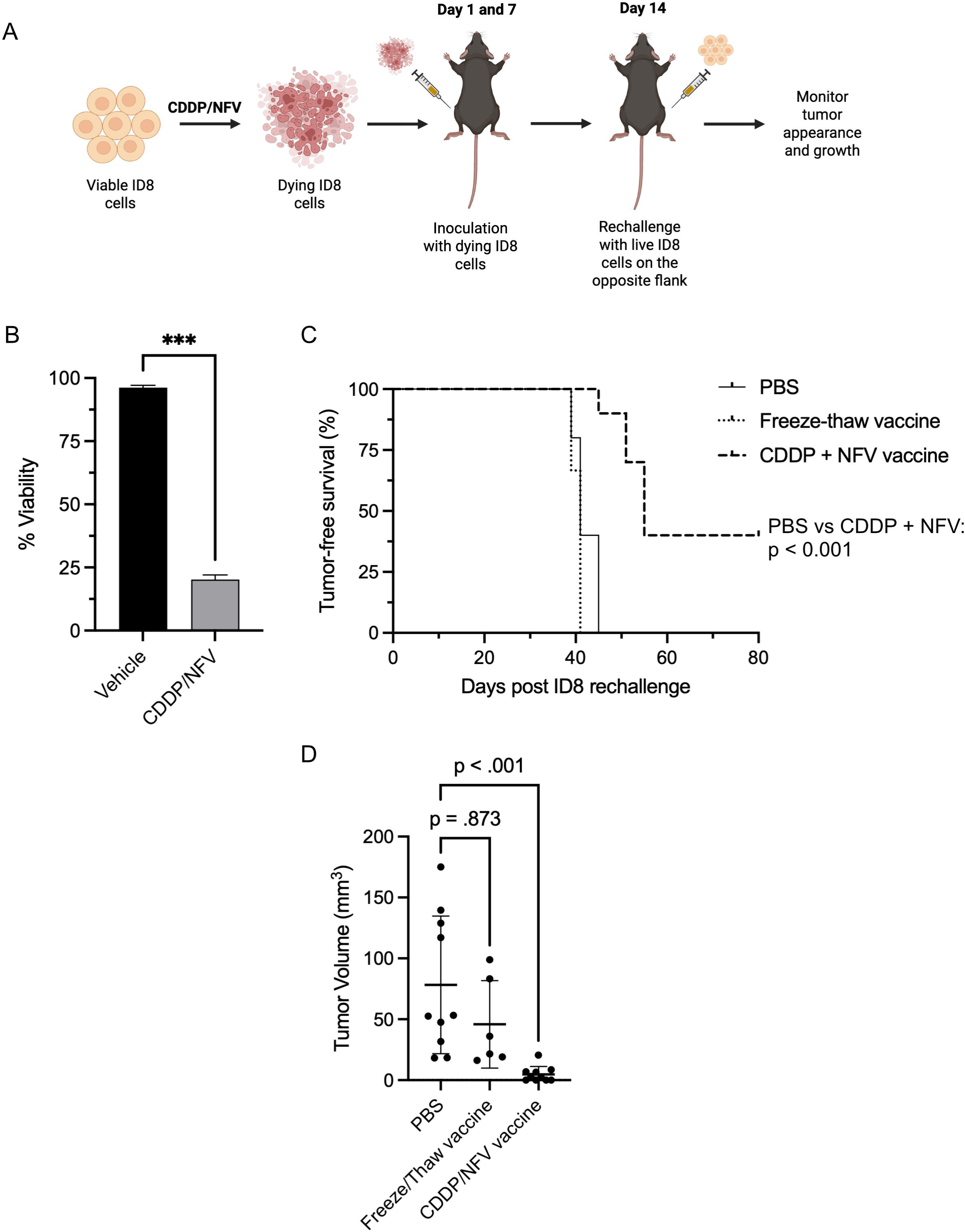
Prophylactic vaccination with dying/dead CDDP/NFV-treated ID8 cells reduces tumor incidence and tumor burden following rechallenge. A) Vaccination model: female C57BL/6 mice received subcutaneous injections of dying CDDP/NFV-treated ID8 cells on Days 1 and 7, followed by contralateral rechallenge with live untreated ID8 cells on Day 14. Tumor development was monitored. B) ID8 cell viability following vehicle or CDDP/NFV treatment before vaccine preparation. Data are presented as mean ± SD from three independent biological experiments. \*\*\**p* < 0.001 by unpaired two-tailed Student’s *t*-test. C) Kaplan–Meier analysis of tumor-free survival (PBS, n = 10; freeze–thaw vaccine, n = 6; CDDP + NFV vaccine, n = 10). Tumor onset was defined by first palpable detection. Four mice in the CDDP + NFV vaccine group remained tumor-free and were censored at day 80. Survival did not differ between PBS and freeze-thaw groups (p = 0.12) but improved significantly with CDDP + NFV vaccination versus PBS (p < 0.001). D) Endpoint tumor volumes; each point represents one mouse, with tumor-free mice assigned 0 mm³. Horizontal lines and error bars indicate mean ± SD. Statistical analysis used the Kruskal–Wallis test followed by Dunn’s multiple-comparisons test.

CDDP/NFV treatment markedly reduced ID8 cell viability relative to vehicle treatment, confirming that the vaccine preparation consisted predominantly of dying tumor cells and demonstrating that the combination was also highly cytotoxic to murine ID8 cells, as previously observed in human OC cells (Fig. 4B). As a non-ICD control, a separate group of mice received a vaccine preparation of ID8 cells killed by accidental necrosis following two consecutive freeze-thaw cycles. This form of cell death is not generally associated with the release of adjuvants and antigens in quantities sufficient to promote ICD [17,43]. Pairwise Mantel-Cox log-rank comparisons showed no significant difference in tumor-free survival between the PBS and freeze-thaw vaccine groups (*p* = 0.12; Fig. 4C). In contrast, tumor-free survival was significantly prolonged in the CDDP/NFV vaccine group relative to the PBS controls (*p* < 0.001). All mice in the PBS group developed palpable tumors by day 45 after rechallenge, as did all mice in the free-thaw vaccine group by day 41. In contrast, tumor development was delayed in the CDDP/NFV vaccine group, while four of ten mice remained tumor-free through the day 80 study endpoint.

Consistent with the improvement in tumor-free survival, endpoint tumor volumes among mice that developed tumors were significantly lower in the CDDP/NFV-vaccinated group than in PBS controls (Dunn’s adjusted *p* < 0.001; Fig. 4D). By contrast, endpoint tumor volumes did not differ significantly between the freeze-thaw and PBS groups (Dunn’s adjusted *p* = 0.873). Together, these findings demonstrate that vaccination with CDDP/NFV-treated ID8 cells provides functional protection against subsequent live tumor challenge, supporting the immunogenicity of CDDP/NFV-induced cell death.

## 3. Discussion

In this study, we show that NFV redirects CDDP-induced OC cell death from a reportedly non-inflammatory phenotype [22] toward an immunogenic phenotype associated with cytoplasmic dsDNA accumulation, STING pathway activation, inflammatory transcriptional reprogramming, extracellular release of soluble mediators, and protective anti-tumor immunity in a prophylactic vaccination model. Whereas CDDP alone did not strongly activate the immune-associated readouts examined, NFV alone and combined with CDDP induced IFN-associated transcriptional programs, increased chemokine and antigen-presentation gene expression, promoted cytoplasmic dsDNA accumulation, and stimulated the release of mediators associated with immunogenic and inflammatory cell death [17,19]. Thus, NFV may alter not only the extent of CDDP-induced OC cell death we reported [32], but also its immunological consequences.

A central finding was that NFV induced transcriptional programs associated with IFN signaling, cytoplasmic DNA sensing, chemokine production, and class I antigen presentation, alone or combined with CDDP. Effective antitumor immunity requires both antigenicity and adjuvanticity: tumor antigens must be available for immune recognition, while dying cells must provide inflammatory signals that support antigen-presenting cell activation and adaptive immune priming [17–19,44]. Increased expression of *CXCL10*, *CXCL11*, *CCL5*, and other inflammatory chemokines may promote immune cell recruitment into the TIME [45]. Similarly, increased *HLA class 1*, *B2M*, *TAP1*, *TAP2*, and *TAPBP* expression suggests enhanced transcriptional features associated with antigen processing and presentation [46,47]. Conversely, reduced expression of selected immunomodulatory or suppressive transcripts may indicate a partial shift away from an immune-evasive state. Nevertheless, these transcriptional changes do not establish enhanced antigen presentation or immune recognition and require functional validation in tumor-immune cell systems.

Our data further implicate cytoplasmic DNA accumulation as a potential link between NFV-induced cellular stress and inflammatory signaling. NFV, alone or combined with CDDP, reduced mature Lamin A abundance and increased extranuclear dsDNA staining, while ddPCR confirmed increased cytoplasmic *TP53* DNA copies following combination treatment. These findings suggest that altered nuclear-envelope integrity and DNA damage-associated stress may facilitate cytoplasmic accumulation of self-DNA [48,49]. In IGROV-1/CP and TOV21G cells, these changes coincided with STING and TBK1 phosphorylation, supporting activation of cytoplasmic DNA-associated inflammatory signaling [48,50]. However, downstream STING activation varied in PEO4 cells despite cytoplasmic dsDNA accumulation, suggesting these events may not be concordant across OC cell lines or histotypes. Differences in pathway competency, activation kinetics, negative feedback, or signaling thresholds may account for this variability. Time-course studies and genetic or pharmacologic cGAS/STING modulation are needed to establish causality. Furthermore, other cytoplasmic dsDNA sensors, including the AIM2 inflammasome [51] should be investigated, particularly in cells lacking detectable STING pathway activation.

Consistent with these transcriptional changes, CDDP/NFV increased the release of soluble mediators associated with inflammation and ICD [17]. HMGB1 release was greatest following combination treatment, whereas IL-18 increased following NFV alone and further with CDDP/NFV. IFN-β was detected in IGROV-1/CP cells treated with NFV alone or the combination, but not in PEO4 cells, consistent with cell-line-dependent STING-TBK1 activation.

These mediators have distinct immune functions: HMGB1 acts as a DAMP [37,52], IL-18 promotes inflammatory immune activation [53], and type I IFNs support dendritic cell function, antigen presentation, and effector lymphocyte responses [54]. Importantly, ICD requires more than DAMP release; dying cells must provide antigenic and adjuvant signals capable of initiating antitumor immunity *in vivo* [17,55]. Accordingly, the combined release of DAMPs, inflammatory cytokines, and type I IFN-associated signals, together with tumor protection in the prophylactic vaccination assay, supports the immunogenic nature of CDDP/NFV-induced OC cell death and suggests stronger antitumor immune potential than Pt-based chemotherapy alone.

The strongest functional evidence came from the prophylactic vaccination model. Vaccination with CDDP/NFV-treated mouse OC ID8 cells reduced tumor incidence and endpoint tumor burden following live tumor rechallenge, whereas freeze-thaw-killed cells provided no comparable protection. Freeze-thaw killing controls for non-programmed or poorly immunogenic cell death associated with accidental necrosis and lacking coordinated antigenicity and adjuvanticity [43]. Thus, the protection afforded by CDDP/NFV-treated cells suggests immune-stimulatory properties beyond passive release of intracellular contents. Although additional immune profiling is needed to identify the responsible effector mechanisms, reduced tumor incidence and burden support the induction of protective antitumor immunity [17].

This study has several limitations. First, we performed RNA-seq only in IGROV-1/CP cells, although we examined key downstream phenotypes in PEO4 and TOV21G cells. Second, although we observed an association between NFV exposure, cytoplasmic DNA accumulation, STING-TBK1 activation, and inflammatory signaling, we did not directly establish whether cGAS/STING is required. Third, STING-associated signaling was more evident in IGROV-1/CP and TOV21G than in PEO4 cells. These models differ in histological origin: TOV21G was classified as differentiated endometrioid OC by transcriptional profiling [56,57], PEO4 was derived from a patient with HGSOC [58], and the parental IGROV-1 line originated from a tumor with mixed histological features [59], although it clusters transcriptionally with ovarian clear-cell carcinomas [56]. STING expression also varies among OC histotypes, although its relationship with treatment-induced pathway activation remains unclear [60]. Thus, histotype-associated biology and other model-specific characteristics may contribute to differential STING pathway responses. Finally, although the vaccination experiment demonstrates functional immunogenicity, the immune cell populations mediating protection remain undefined.

Future studies should investigate dendritic cell activation, antigen-specific T-cell responses, immune memory, and the therapeutic potential of combining CDDP/NFV with immune-based strategies to induce ICD *in vivo*. By promoting ICD, this combination may potentially convert immunologically “cold” ovarian tumors into “hot” tumors more responsive to ICIs, which currently provide limited clinical benefit in most OC patients [7].

In summary, NFV shifts CDDP-induced OC cell death toward an immunogenic phenotype characterized by cytoplasmic DNA accumulation, STING-associated inflammatory signaling, DAMP and cytokine release, and protective anti-tumor immunity. These findings support further investigation of NFV as a repurposed agent that can enhance both the cytotoxic and immunogenic consequences of Pt-based therapy in OC.

## CrediT authorship contribution statement

**Benjamin N. Forgie:** Conceptualization, Writing-original draft, Methodology, Investigation, Formal analysis, Visualization. **Sarah T. Ferrier:** Methodology, Investigation, Formal analysis, Review and Editing. **Alicia A. Goyeneche:** Supervision, Methodology, Investigation, Review. **Alexandra Bartolomucci:** Methodology, Review and Editing. **Desiree Marno:** Investigation, Review and Editing. **Rewati Prakash:** Formal analysis; **Edith Zorychta:** Review and Editing. **Abu Shadat M. Noman:** Funding acquisition, Review. **Lucy Gilbert:** Funding acquisition, Review. **Julia V. Burnier:** Writing-Review and Editing, Supervision, Funding acquisition, Resources. **Carlos M. Telleria:** Writing-Review and Editing, Supervision, Resources, Project Administration, Funding Acquisition, Conceptualization.

## Declarations of competing interests

The authors declare that they have no known competing financial interests or personal relationships that could have appeared to influence the work reported in this paper.

## Acknowledgements

This work was supported by grants from DxQuest Inc. North America and Ovarian Cancer Canada (to C.M.T.), internal funds from the Gerald Bronfman Department of Oncology at McGill University (to L.G.), and a Canada Graduate Scholarship Doctoral Research Award (CGSD) (to B.N.F.; Award #193306). J.V.B. was supported by Fonds de Recherche du Québec en Santé (Award #312831).

## Data availability

The RNA-seq data analyzed in this study are publicly available through the Gene Expression Omnibus (GEO) under accession number GSE338229. Other data supporting the findings are available from the corresponding author upon reasonable request.

## Abbreviations

OC: Ovarian cancer
NFV: Nelvinafir
CDDP: Cisplatin
Pt: Platinum
OXP: Oxaliplatin
CP: Carboplatin
ICD: Immunogenic cell death
TIME: Tumor immune microenvironment
ICIs: Immune checkpoint inhibitors
DAMPs: Damage-associated molecular patterns
IFN: Interferon
IL-18: Interleukin-18
STING: Cyclic GMP–AMP synthase (cGAS)–stimulator of interferon genes
TBK1: TANK-binding kinase 1
PD-L1: Programmed Death-Ligand 1
dsDNA: Double-stranded DNA
ddPCR: Droplet digital polymerase chain reaction

---

**Suppl. Fig. 1.**
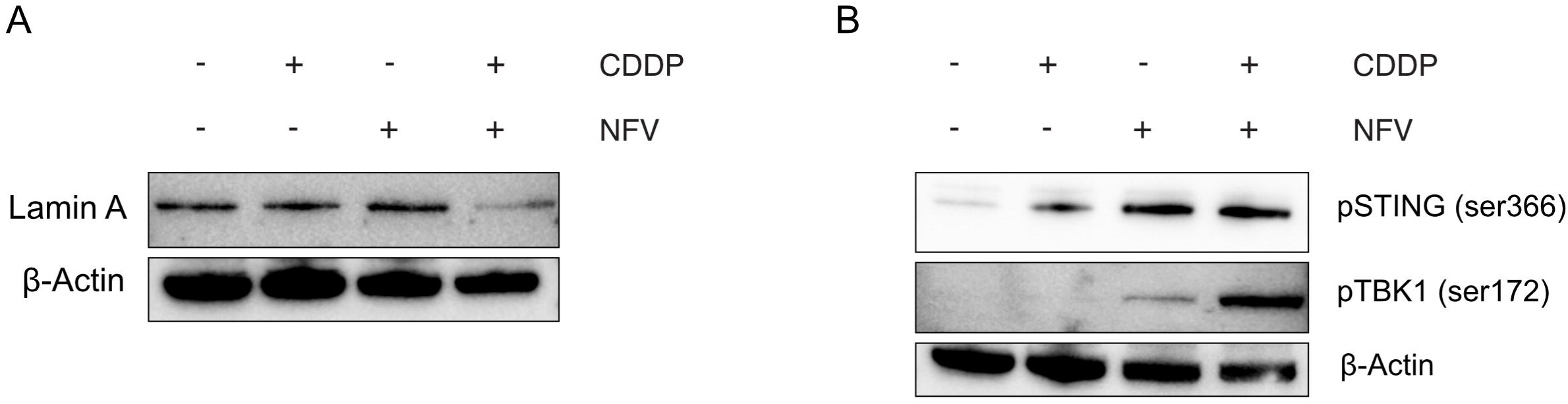
CDDP/NFV treatment reduces lamin A abundance and promotes STING-pathway phosphorylation in TOV21G cells. A) Representative immunoblot showing Lamin A levels following vehicle, CDDP, NFV, or CDDP + NFV. B) Representative immunoblots showing STING phosphorylation (ser366) and TBK1 phosphorylation (ser172) following the indicated treatments. β-Actin served as a loading control. Immunoblots are representative of two independent experiments.

## Notes

### Competing Interest Statement

The authors have declared no competing interest.

